# Deaminase associated single nucleotide variants in blood and saliva-derived exomes from healthy subjects

**DOI:** 10.1101/807073

**Authors:** Nathan E. Hall, Jared Mamrot, Christopher M.A. Frampton, Prue Read, Edward J. Steele, Robert J. Bischof, Robyn A. Lindley

## Abstract

**Background:** Deaminases play an important role in shaping inherited and somatic variants. Disease related SNVs are associated with deaminase mutagenesis and genome instability. Here, we investigate the reproducibility and variance of whole exome SNV calls in blood and saliva of healthy subjects and analyze variants associated with AID, ADAR, APOBEC3G and APOBEC3B deaminase sequence motifs.

**Methods:** Samples from twenty-four healthy Caucasian volunteers, allocated into two groups, underwent whole exome sequencing. Group 1 (n=12) analysis involved one blood and four saliva replicates. A single saliva sample was sequenced for Group 2 subjects (n=12). Overall, a total of 72 whole exome datasets were analyzed. Biological (Group 1 & 2) and technical (Group 1) variance of SNV calls and deaminase metrics were calculated and analyzed using intraclass correlation coefficients. Candidate somatic SNVs were identified and evaluated.

**Results:** We report high blood-saliva concordance in germline SNVs from whole exome sequencing. Concordant SNVs, found in all subject replicates, accounted for 97% of SNVs located within the protein coding sequence of genes. Discordant SNVs have a 30% overlap with variants that fail gnomAD quality filters and are less likely to be found in dbSNP. SNV calls and deaminase-associated metrics were found to be reproducible and robust (intraclass correlation coefficients >0.95). No somatic SNVs were conclusively identified when comparing blood and saliva samples.

**Conclusions:** Saliva and blood both provide high quality sources of DNA for whole exome sequencing, with no difference in ability to resolve SNVs and deaminase-associated metrics. We did not identify somatic SNVs when comparing blood and saliva of healthy individuals, and we conclude that more specialized investigative methods are required to comprehensively assess the impact of deaminase activity on genome stability in healthy individuals.

## Background

APOBEC/AID deaminases are a recognized endogenous source of genome instability [1–5]. Somatic mutations caused by deamination events have been identified in cancer *in vitro* and *in vivo* [6–9], and evidence of deaminase-associated mutations in non-cancerous conditions is emerging, such as various viral infections and neurodegenerative diseases [10,11]. Deaminases have also recently been implicated in accumulation of pre-cancerous mutations [12], and as a causative driver of many human SNPs [13].

Deaminases predominantly drive C-to-U(T) and A-to-I(G) transition mutations, however DNA repair mechanisms typically prevent deamination from compromising genome integrity and causing somatic mutation [14,15]. Pathophysiological processes can disrupt normal DNA repair, resulting in mosaic manifestation of deaminase-associated single nucleotide variants (SNVs) in affected tissues [16]. Although deaminases employ similar biochemical mechanisms, each has a unique binding domain associated with one or more DNA motifs [17,18]. Deaminase motifs can be identified and quantified in Next-Generation Sequencing (NGS) data facilitating diagnosis of the specific cause of the mutation. For example, AID targets C-sites in the context of WR<u>C</u> motifs (W = A or T; R = A or G; reverse complements as <u>G</u>YW, with Y = T or C), APOBEC3G deaminates C<u>C</u> sites (or <u>G</u>G) and APOBEC3B deaminates T<u>C</u>W (or W<u>G</u>A) motifs and ADARs deaminate W<u>A</u> sites [2,19,20]. Establishing reproducible and robust deaminase-associated SNV profiles in healthy people will improve the utility of mutation profiling techniques for monitoring progression of diseases such as cancer, and for understanding patient response to treatment.

Sampling of saliva or buccal cells is a widely employed technique for collecting human DNA for ancestry, forensic, medical and research purposes [21,22,23,24]. DNA extracted from saliva can be analyzed using various NGS techniques, however the quality of DNA derived from saliva can be compromised by metagenomic DNA and activity of various enzymes and antibacterial factors. There are several practical advantages to this DNA source, such as ease of sampling and additional sequencing information about metagenomic populations [25,26,27], however DNA obtained from saliva is not yet routinely used for detecting SNVs.

Here, we report profiles for SNVs associated with deaminase motifs for a cohort of 24 healthy human subjects using whole exome sequencing (WES). For twelve of these subjects (Group 1) we compare blood with biological and technical saliva replicates from Caucasian volunteers of different age groups and sex and hypothesize that deaminase-associated SNV profiles of a cohort of healthy individuals will show a high concordance between saliva and whole blood DNA in a reproducible and robust manner.

## Methods

### Healthy subject selection

In total, 24 healthy Caucasian subjects were recruited for this study. Volunteers were considered healthy if they had blood pressure and heart rate within normal ranges, had never smoked, were only light drinkers (<14 units of alcohol weekly), had no major viral infections or immune related diseases and did not take any regular medication. Eight subjects were recruited into each of the three age groups 18-19, 30-39, and 50-59, with an equal ratio of males to females in each group. These subjects were randomly allocated into two groups of equal sex and age group. Group 1 (n=12) involved analysis of blood and saliva sample replicates. Group 2 (n=12) involved analysis of saliva-1 sample only. This project was approved by the Monash Health Human Research Ethics Committee (16281L: “A study to measure the Targeted Somatic Mutation (TSM) test platform performance characteristics and evaluate its suitability for clinical use”).

### Sample collection

For each subject, two saliva samples were collected, 30 minutes apart, using the Oragene DNA (OG-500) saliva collection kit. Whole blood samples were collected into sterile EDTA tubes.

### DNA extraction from saliva and whole blood

DNA was extracted from samples using the QIAsymphony and the Qiagen DSP DNA Mini Kit. The extracted DNA was eluted in 100uL of Qiagen ATE buffer.

### Library preparation

Whole exome sequencing library preparation was performed at the Monash Health Translation Precinct (MHTP) Medical Genomics Facility using the Agilent SureSelectXT Target Enrichment System according to protocol G7530-90000, Version C0, December 2016. Capture Probes: Agilent SureSelect Clinical Research Exome Cat No 5190-7344; Design ID S06588914. Libraries were QC-checked using the Agilent BioAnalyzer and quantified with Qubit.

### Whole Exome Sequencing

Four samples per lane were clustered on the c-bot using 200pM of library pool using Illumina Protocol 15006165 v02 Jan 2016. Raw data was generated on the Illumina HiSeq 3000 with 100 base-pair paired-end (PE) sequencing with Illumina Protocol 15066493 Rev A, February 2015. Total PE reads per sample were between 110 million and 183 million per exome, excluding HP_4 saliva-1 which had 82 million. The median number of PE reads per sample (137 million) and additional summary statistics are provided in Supplementary Table 1.

### Bioinformatics analysis

WES read quality was assessed using FastQC (v0.11.7) [28]. Adapters were trimmed with cutadapt (v1.16) [29] and mapped to the human genome version hg19 with bwa (v0.7.13-r1126) [30] with the parameters “bwa mem -M -t 5 -k 19”. Duplicates were marked with Picard MarkDuplicates (v2.6.0) (http://broadinstitute.github.io/picard). Single Nucleotide Variant (SNV) calls were made with Strelka2 (v2.8.4) [31] with default parameters using “configureStrelkaGermlineWorkflow.py -exome”. Variants failed quality filtering if they had a ConservativeGenotypeQuality < 15, a RelativeTotalLocusDepth < 3, or a SampleStrandBias > 10. Variants remaining after quality filtering were converted from hg19 to hg38 coordinates using UCSC’s LiftOver tool [32]. Variants were identified as being located in the coding sequence (CDS) of genes according to Ensembl version 92 [33]. Candidate SNVs were compared against dbSNP v150 (10-07-2017) [34] and gnomAD exome release v2.0.2 [35]. Candidate somatic SNVs were identified using Strelka2 with default parameters in the “configureStrelkaSomaticWorkflow.py -exome” pipeline. Unmapped reads were QC checked using FastQC and MultiQC (v1.6) [36] (https://jpmam1.github.io/MultiQC). To determine the source of the unmapped reads from representative subjects HP_1 and HP_2, these were aligned to the NCBI “non-redundant” (nr) database comprised of 4,348,972 protein sequences from eukaryotic and prokaryotic organisms, using DIAMOND BLASTx (v0.9.22.123) [37]. Alignments were visualized using MEGAN6 (v6.12.2) [38,39].

### Deaminase motifs in WES data

SNVs occurring within four key deaminase motifs were identified and quantified. The motifs used were AID: WR<u>C</u> / <u>G</u>YW, ADAR: W<u>A</u> / <u>T</u>W, APOBEC3G: C<u>C</u> / <u>G</u>G, APOBEC3B: T<u>C</u>W / W<u>G</u>A where W=A or T, Y=C or T, R=A or G [2,19,20]. The base mutated in each motif is underlined. Motif searches were conducted according to the direction of the gene. Transition/transversion ratios (Ti/Tv) were calculated as the proportion of total transition variants. Strand bias was calculated as the proportion of variants on the forward strand (e.g. C:G and A:T as percentages). Motif-independent metrics and SNVs not associated with motifs of AID, ADAR, APOBEC3G or APOBEC3B (denoted “Other”) were also quantified.

### Experimental design

Five WES datasets were generated for twelve subjects (Group 1), comprised of two males and two females from three age categories (18-19, 30-39 and 50-59). As described in Figure 1, replicates were generated from two saliva samples at the DNA extraction stage (saliva-1C) and at the library preparation stage (saliva-2A). These technical and biological replicates enabled analysis of concordance between replicates and provided a measure of technical variance and noise. This study design allows quantitative comparisons between blood and saliva, between saliva sample replicates and between technical saliva replicates at the DNA extraction and library preparation level. Group 2 subjects (n=12) underwent WES of saliva-1 samples only and were used in the calculation of biological variance between subjects.

**Figure 1.**
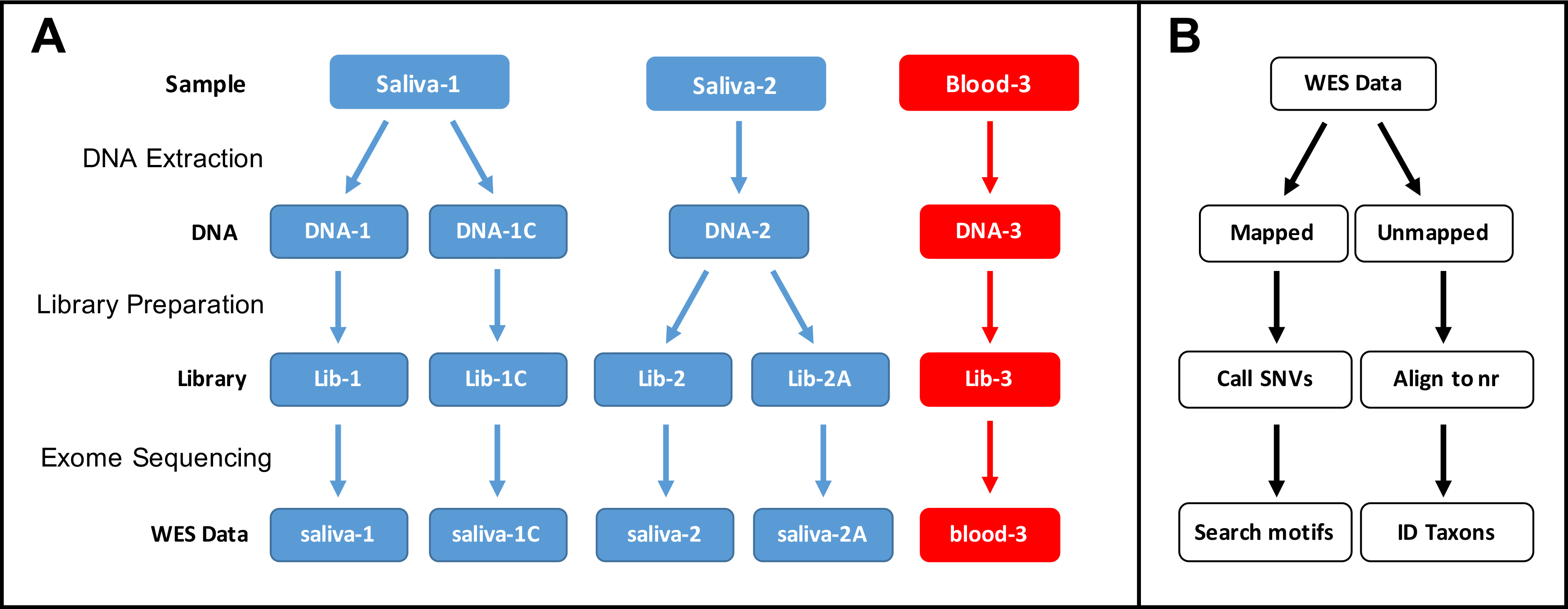
Experimental design. (**A**) Flow diagram of whole exome sequencing (WES) pipeline. DNA was extracted from two saliva samples “1” and “2” and a blood sample “3” to make libraries for whole exome sequencing. Lib-1C is made from a separate DNA extraction of Sample 1, and Lib-2A is a separate library made from the same Saliva-2 DNA extraction. Blood samples have only one DNA extraction and one library preparation. (**B**) WES data processing pipeline. Reads were aligned to the reference human genome, SNVs were called in mapped reads and variants associated with deaminase motifs were quantified. Unmapped reads were aligned to the NCBI ‘non-redundant’ (nr) to establish the taxonomic sources of reads.

Mapped and unmapped WES reads were analyzed for genomic variants and off-target metagenomic contamination.

### Statistical analyses

Intraclass correlation was calculated for SNV counts and deaminase motif metrics using the formula described by Shrout & Fleiss [40]: 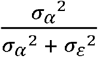. In brief, σ_α_ represents the biological variance between subjects and σ_ε_ represents the technical variance within subject replicates. Used here, σ_α_ is the standard deviation of saliva-1 samples across all 24 subjects (Group 1 and 2). For each Group 1 subject (n=12), a standard deviation is calculated and represents the variance within blood and saliva samples, DNA extraction and libraries; σ_ε_ is the average of these twelve standard deviations.

SNVs termed *discordant* were found in 1, 2, 3, or 4 of the 5 samples, but not in all samples for each individual. Venn diagrams of concordant/discordant SNVs were generated at http://bioinformatics.psb.ugent.be/webtools/Venn/. Pairwise sample comparisons were conducted for discordant SNVs and analyzed using one-way ANOVA.

## Results

### Blood and saliva whole exome sequencing

Saliva and blood samples from 12 healthy volunteers, Group 1 subjects, underwent sequencing and analysis according to the workflow illustrated in Figure 1. In addition, 12 exomes were obtained from Saliva-1 samples from the remaining 12 recruited healthy volunteers, Group 2 subjects (Table 2). For all exomes sequenced (n=72), an average of 136 million high-quality 100bp paired-end reads were obtained. The total number of reads, mapping rate and coverage statistics for all sequencing runs are described in Supplementary Table 1. Mapping rates were between 94.2% and 99.9% with a median of 98.9%. The median exome coverage rates were 97.2% (>30×) and 70.0% (>100×) of the exome. Sample HP_4_1 produced the lowest number of reads and subsequently had the lowest sequencing depth with 91.5% of the exome covered by >30×. Age group, sex, and counts for total SNVs, SNVs within a coding region (referred to CDS), and percentages of variants within a coding sequence region that correspond to known motifs for AID, ADAR, APOBEC3G and APOBEC3B are presented in Tables 1 and 2, and Supplementary Figure 3.

**Table 1:** Summary of total SNV counts, CDS SNV counts, and percentages of CDS variants that correspond to motifs for AID, ADAR, APOBEC3G and APOBEC3B for Group 1 subjects, comprising one blood and four saliva replicate datasets. Sex and age group is given for each healthy subject.

| ID |  | HP_15 | HP_16 | HP_3 | HP_10 | HP_1 | HP_4 | HP_12 | HP_20 | HP_2 | HP_5 | HP_6 | HP_22 |
| --- | --- | --- | --- | --- | --- | --- | --- | --- | --- | --- | --- | --- | --- |
| Sex |  | M | M | F | F | M | M | F | F | M | M | F | F |
| Age Group |  | 18-19 | 18-19 | 18-19 | 18-19 | 30-39 | 30-39 | 30-39 | 30-39 | 50-59 | 50-59 | 50-59 | 50-59 |
| SNV | saliva-1 | 44227 | 44334 | 44913 | 44075 | 44411 | 44249 | 44531 | 44848 | 44121 | 44389 | 44256 | 44306 |
| SNV | saliva-1C | 44261 | 44408 | 45008 | 44196 | 44366 | 44354 | 44637 | 44914 | 44218 | 44444 | 44423 | 44419 |
| SNV | saliva-2 | 44346 | 44509 | 44910 | 44222 | 44387 | 44291 | 44609 | 45051 | 44133 | 44304 | 44391 | 44356 |
| SNV | saliva-2A | 44305 | 44459 | 44934 | 44241 | 44436 | 44354 | 44616 | 44974 | 44170 | 44372 | 44414 | 44430 |
| SNV | blood-3 | 44247 | 44338 | 44890 | 44124 | 44479 | 44297 | 44520 | 44963 | 44073 | 44340 | 44398 | 44324 |
| CDS | saliva-1 | 20236 | 20309 | 20649 | 20084 | 20358 | 20326 | 20399 | 20605 | 20245 | 20359 | 20054 | 20233 |
| CDS | saliva-1C | 20280 | 20369 | 20651 | 20123 | 20358 | 20308 | 20418 | 20663 | 20306 | 20345 | 20131 | 20250 |
| CDS | saliva-2 | 20293 | 20379 | 20641 | 20121 | 20380 | 20283 | 20408 | 20713 | 20275 | 20313 | 20136 | 20256 |
| CDS | saliva-2A | 20268 | 20398 | 20634 | 20126 | 20373 | 20298 | 20427 | 20655 | 20301 | 20327 | 20148 | 20277 |
| CDS | blood-3 | 20255 | 20305 | 20620 | 20127 | 20415 | 20289 | 20389 | 20646 | 20255 | 20301 | 20136 | 20218 |
| AID% | saliva-1 | 13.72 | 13.71 | 13.80 | 13.63 | 14.03 | 13.69 | 13.68 | 13.63 | 13.63 | 13.56 | 13.60 | 13.88 |
| AID% | saliva-1C | 13.67 | 13.71 | 13.82 | 13.64 | 14.00 | 13.67 | 13.67 | 13.69 | 13.66 | 13.62 | 13.59 | 13.84 |
| AID% | saliva-2 | 13.73 | 13.72 | 13.83 | 13.66 | 14.03 | 13.69 | 13.69 | 13.66 | 13.68 | 13.57 | 13.61 | 13.88 |
| AID% | saliva-2A | 13.69 | 13.72 | 13.79 | 13.66 | 14.02 | 13.69 | 13.65 | 13.63 | 13.62 | 13.60 | 13.63 | 13.86 |
| AID% | blood-3 | 13.72 | 13.69 | 13.81 | 13.60 | 14.02 | 13.70 | 13.68 | 13.62 | 13.68 | 13.55 | 13.60 | 13.90 |
| ADAR% | saliva-1 | 15.40 | 15.43 | 15.67 | 15.85 | 15.57 | 15.54 | 15.45 | 15.64 | 15.56 | 15.66 | 15.45 | 15.64 |
| ADAR% | saliva-1C | 15.43 | 15.43 | 15.67 | 15.83 | 15.59 | 15.50 | 15.46 | 15.65 | 15.54 | 15.64 | 15.45 | 15.64 |
| ADAR% | saliva-2 | 15.37 | 15.41 | 15.66 | 15.83 | 15.58 | 15.48 | 15.47 | 15.64 | 15.60 | 15.67 | 15.47 | 15.61 |
| ADAR% | saliva-2A | 15.42 | 15.39 | 15.69 | 15.81 | 15.59 | 15.50 | 15.46 | 15.62 | 15.57 | 15.66 | 15.47 | 15.58 |
| ADAR% | blood-3 | 15.37 | 15.47 | 15.69 | 15.85 | 15.52 | 15.49 | 15.48 | 15.67 | 15.60 | 15.69 | 15.48 | 15.59 |
| APOBEC3G% | saliva-1 | 16.95 | 16.71 | 16.90 | 16.68 | 16.96 | 16.92 | 17.13 | 16.90 | 16.76 | 16.90 | 16.74 | 16.65 |
| APOBEC3G% | saliva-1C | 16.94 | 16.75 | 16.89 | 16.72 | 16.91 | 16.92 | 17.16 | 16.85 | 16.77 | 16.90 | 16.71 | 16.65 |
| APOBEC3G% | saliva-2 | 16.95 | 16.71 | 16.90 | 16.71 | 16.91 | 16.97 | 17.12 | 16.85 | 16.69 | 16.92 | 16.72 | 16.58 |
| APOBEC3G% | saliva-2A | 16.97 | 16.70 | 16.88 | 16.67 | 16.93 | 16.94 | 17.12 | 16.91 | 16.78 | 16.92 | 16.70 | 16.64 |
| APOBEC3G% | blood-3 | 17.01 | 16.69 | 16.86 | 16.74 | 16.95 | 17.03 | 17.11 | 16.85 | 16.75 | 16.92 | 16.73 | 16.65 |
| APOBEC3B% | saliva-1 | 4.06 | 4.13 | 3.99 | 4.09 | 4.13 | 4.02 | 4.17 | 4.06 | 3.99 | 4.13 | 4.26 | 4.14 |
| APOBEC3B% | saliva-1C | 4.05 | 4.12 | 4.00 | 4.11 | 4.13 | 4.03 | 4.12 | 4.03 | 3.98 | 4.10 | 4.26 | 4.13 |
| APOBEC3B% | saliva-2 | 4.07 | 4.10 | 4.00 | 4.10 | 4.11 | 4.02 | 4.14 | 4.03 | 3.99 | 4.11 | 4.26 | 4.14 |
| APOBEC3B% | saliva-2A | 4.05 | 4.12 | 4.00 | 4.10 | 4.11 | 4.02 | 4.15 | 4.03 | 3.99 | 4.13 | 4.26 | 4.12 |
| APOBEC3B% | blood-3 | 4.05 | 4.11 | 4.00 | 4.10 | 4.13 | 4.00 | 4.14 | 4.03 | 3.99 | 4.13 | 4.26 | 4.14 |

**Table 2:**
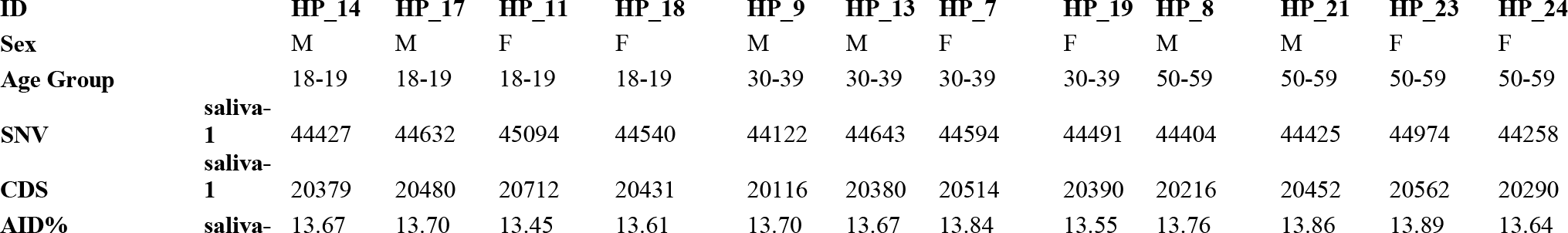
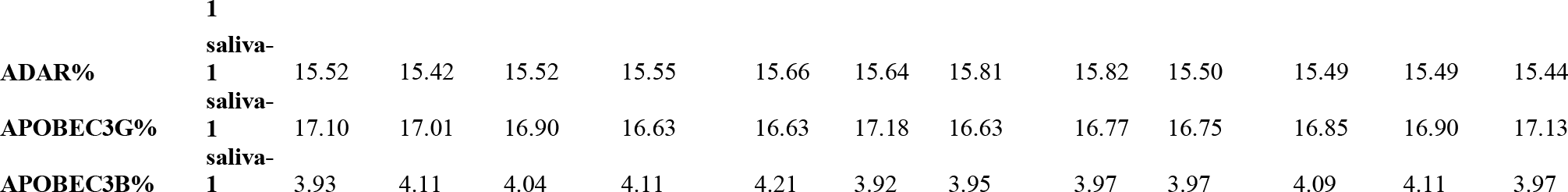
Summary of total SNV counts, CDS SNV counts, and percentages of CDS variants that correspond to the motifs for AID, ADAR, APOBEC3G and APOBEC3B for Group 2 subjects, comprising saliva-1 samples. Sex and age group is given for each healthy subject.

### SNV concordance between and within sample types

For Group 1 subjects (n=12), SNVs called in each sample were analyzed following the workflow described in Figure 1. Variants shared between sample types were quantified (i.e. concordance between saliva ‘1’, ‘1C’, ‘2’, ‘2A’, and blood ‘3’), with all sample types showing very high concordance overall. Venn diagrams illustrate overlap of variant calls in all exome regions, as well as those located within gene coding regions (CDS) between sample types and replicates for a representative volunteer (HP_1: Figure 2A and 2C). Overall, 96.1% of total variants in this volunteer were common to all sample replicates and are referred to as *concordant* SNVs. Venn diagrams for all volunteers are provided in Supplementary Figures 1 (all SNVs) and 2 (SNVs restricted to gene CDS). SNVs common to 1, 2, 3, or 4 but not all 5 of the samples are referred to as *discordant* SNVs and were further investigated. The percentage of concordant SNVs was slightly higher on average in the CDS (96.6%), compared to those in all WES regions (95.8%) (Supplementary Table 2). As a measure of pairwise similarity between samples, the number of *discordant* SNVs in common between sample pairs are shown in Figure 2B, WES SNVs, and Figure 2D, CDS SNVs. Pairwise comparisons are categorized as: biological and technical *blood-saliva replicates* (blood-3 & saliva-1, blood-3 & saliva-2, blood-3 & saliva-1C, blood-3 & saliva-2A), biological and technical *saliva replicates* (saliva-1 & saliva-2, saliva-1 & saliva-2A, saliva-1C & saliva-2, saliva-1C & saliva-2A), and *technical saliva replicates* (saliva-1 & saliva-1C, and saliva-2 & saliva-2A). The sample with lowest coverage (HP_4 saliva-1) was associated with lower pairwise overlap of discordant reads, however this difference was ameliorated when analysis was restricted to only the coding region of genes. A statistical analysis of the pairwise number of discordant SNVs in common (WES and CDS SNVs) showed no significant difference between the pairwise comparisons of blood-saliva, saliva-saliva and technical saliva replicates (ANOVA; p>0.05).

**Figure 2:**
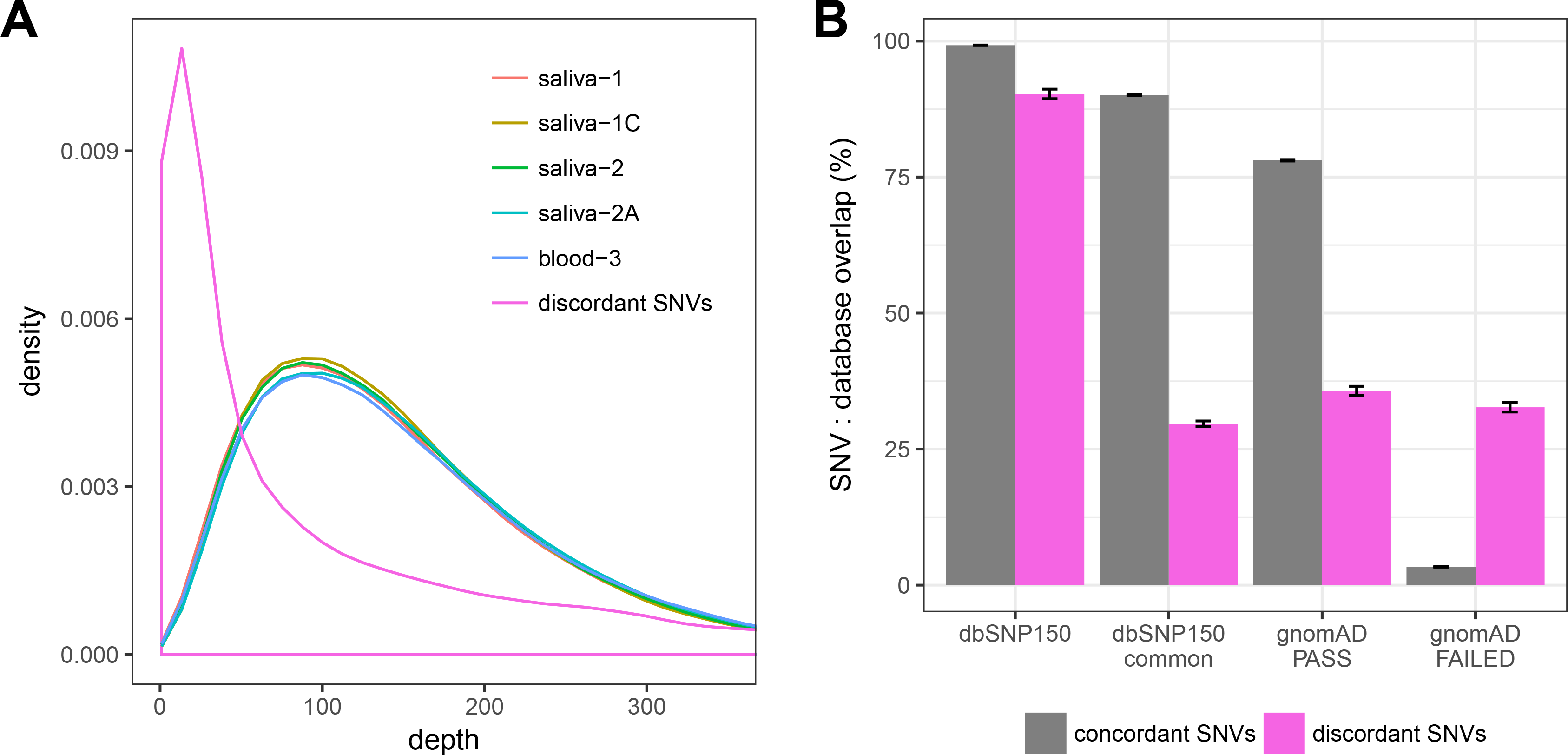
Relationship of SNV calls between and among sample replicates. (**A**) Venn diagram of all called variants by ID, (**B**) pairwise sample comparisons of all discordant SNVs in common for each WES dataset pair, (**C**) Venn diagram of CDS variants, (**D**) pairwise sample comparisons of all discordant CDS SNVs in common for each WES dataset pair. Blood-saliva pairwise comparisons are in shades of red. Saliva-1-saliva-2 comparisons are in shades of blue, and technical saliva replicates are in green.

Sequencing depth for concordant SNVs and discordant SNVs, averaged across 12 samples, is presented in Figure 3. Sequencing coverage distribution typically centered around 100×. Discordant SNVs have a higher density of low WES coverage (<30×). Depth analysis of individual samples are graphed in Supplementary Figure 4. Analysis of HP_1 replicates revealed discordant SNVs failed one or more quality filters due to high strand bias >10, (13% of 1762 discordant SNVs), low genotype quality (62%), high ratio of quality-filtered bases (8%), low depth (7%), or were not called as variants in one or more samples (54%). Overlap of concordant and discordant SNVs with dbSNP and gnomAD databases showed clear differences (Figure 3B). Concordant SNVs have a much higher overlap in dbSNP than discordant SNVs for both ‘all variants’ and ‘common’ variants. Using large-scale analysis of over 120 thousand exomes, the gnomAD database flags variants that do not pass certain quality filters. Of all concordant SNVs, 3% failed the gnomAD filters, however 31% of the discordant SNVs failed.

**Figure 3:**
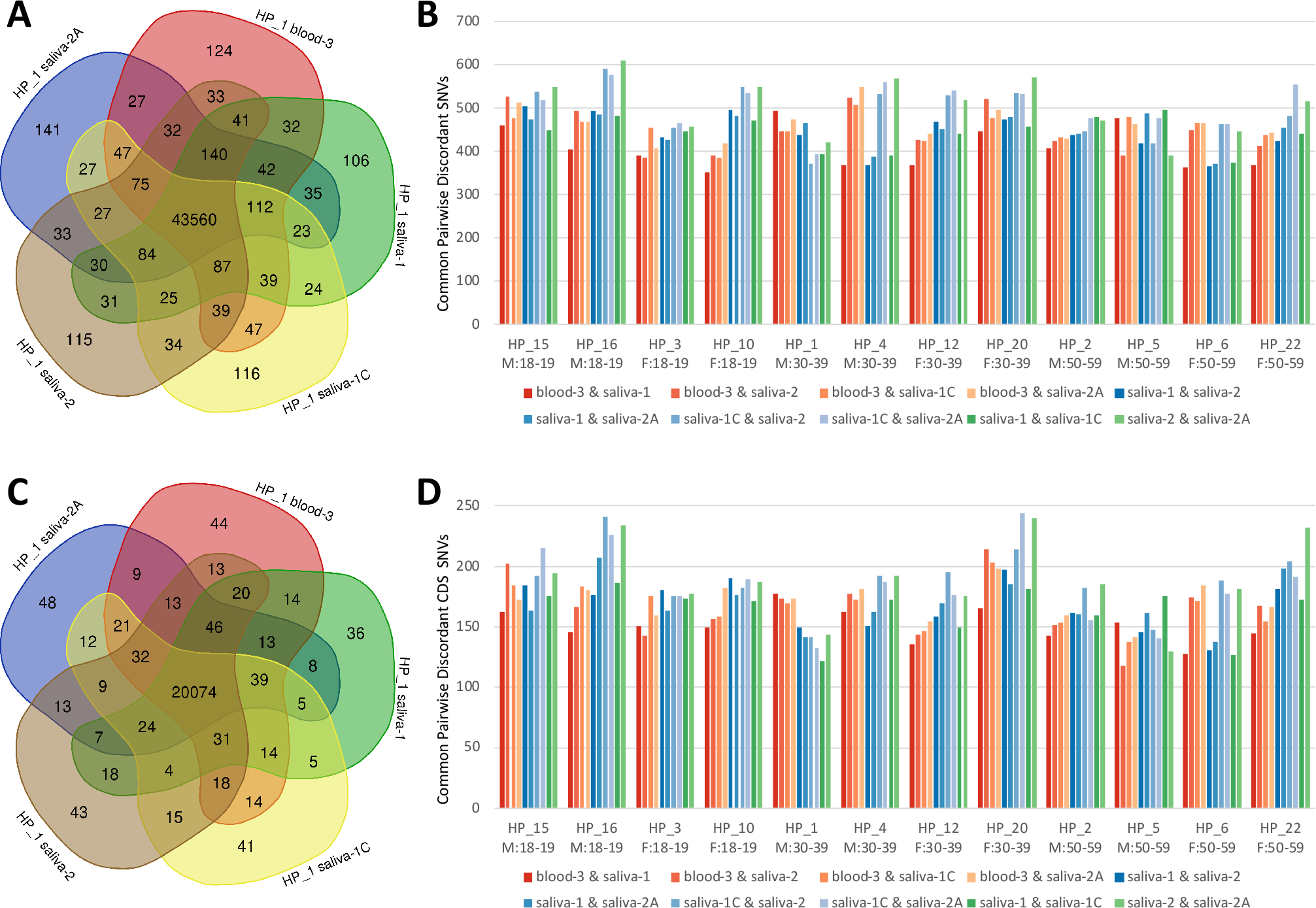
Sequencing depth and database overlap of concordant and discordant SNVs. (**A**) Combined depth profiles of concordant SNVs across five samples types compared to discordant SNVs, and (**B**) the rate of concordant and discordant SNVs belonging to each variant database (n=12, mean ± 95% CI).

### Candidate somatic SNV analysis

Candidate somatic variants were identified using Strelka2 ‘tumor-normal’ methods, with blood and saliva sample replicates alternatingly used as ‘tumor’ and ‘normal’. There was no overlap between discordant SNVs identified in the germline and candidate somatic SNVs. Although all candidate somatic variants passed default filters, the quality of candidate somatic SNVs measured using the Strelka2 Empirical Variant Score (EVS) were all relatively low (EVS < 20). EVS is a phred-scaled probability of the call being a false positive observation and is calculated from pre-trained random forest models and not hard cutoffs[31]. Low EVS scores are typically due to low minor allele frequencies, low mapping quality and low sequence coverage regions[31].

The mean number of somatic SNV candidates found in saliva was 149 (saliva=tumor, blood=normal), those found in blood was 121 (blood=tumor, saliva=normal). There was no correlation detected between the number of candidate somatic variants and the age of subjects. The average number of candidate saliva-blood somatic SNVs was 158, 141 and 148, and blood-saliva averages were 119, 131 and 111 across the age groups 18-19, 30-39 and 50-59 respectively. A measure of technical noise is given by the number of candidate somatic variants found in biological and technical saliva replicates, which were on average 126 and 123 SNVs respectively (Supplementary Figure 5).

Of the candidate somatic SNVs identified, approximately 80% had a variant minor allele frequency <0.05. Applying this conventional filter reduced the average number of candidates per category (saliva, blood, technical replicates, biological replicates) to 34, 31, 23 and 25 respectively. A minimum depth filter of >30 for both ‘tumor’ and ‘normal’ samples further reduced average number of somatic candidates per category to 22, 19, 14 and 15 respectively.

After filtering of the candidate saliva SNVs that were not detected in all saliva replicates, 22 candidates remained with 21 of these found in more than one subject. Manual inspection using IGV suggests a false positive caused by incorrect mapping of soft clipped reads. With only a single blood sample per subject, equivalent filtering of candidate somatic SNVs found only in blood was not possible. The number of candidate somatic SNVs in blood was no larger than the level of technical and biological noise.

### Deaminase associated SNVs

SNVs corresponding to known deaminase motifs (AID: WR<u>C</u> / <u>G</u>YW, ADAR: W<u>A</u> / <u>T</u>W, APOBEC3G: C<u>C</u> / <u>G</u>G, APOBEC3B: T<u>C</u>W / W<u>G</u>A) were identified within the coding region of genes (Tables 1 & 2, Supplementary Figure 3). Deaminase-associated SNVs at the genotype level were highly concordant and similar to the percentage concordance of all CDS SNVs: AID (96.1%), ADAR (97.0%), APOBEC3G (96.3%), APOBEC3B (96.2%) and CDS (96.6%) (Supplementary Table 2). Strand bias and transition/transversion ratios were calculated for each deaminase and are summarized below (Table 3). In total, approximately 50% of the CDS SNVs correspond to AID, ADAR, APOBEC3G and APOBEC3B deaminase motifs. The intraclass correlation coefficient (ICC) was used to quantify the reproducibility of the deaminase metric calculations. The ICC values range between 0.958 and 0.989 for all SNVs, deaminase motifs and associated metrics, revealing a very high consistency/reproducibility among the five samples. ICC calculations for a range of additional metrics are reported in Supplementary Table 3.

**Table 3:** Intraclass correlation coefficients (ICC) illustrating consistency between replicates in relation to SNV calls and the existence of deaminase motifs, transition/transversion metrics (Ti/Tv), and strand bias metrics. The variation *within* Group 1 replicates (n=12, 60 WES datasets) represents a measure of technical reproducibility across blood and saliva samples. Variation *between* samples (n=24 Group 1 and 2 saliva-1 samples) represents a measure of biological variability.

|  | Mean<sup>a</sup> | Variance *within* replicates<sup>b</sup> | Variance *between* individuals<sup>c</sup> | ICC |
| --- | --- | --- | --- | --- |
| Exome SNVs | 44469 | 57.371 | 273.809 | 0.958 |
| CDS SNVs | 20366 | 25.0273 | 170.131 | 0.979 |
| AID % | 13.704 | 0.020 | 0.128 | 0.976 |
| ADAR % | 15.572 | 0.021 | 0.129 | 0.974 |
| APOBEC3G % | 16.862 | 0.026 | 0.169 | 0.978 |
| APOBEC3B % | 4.065 | 0.009 | 0.091 | 0.989 |
| AID Ti/Tv % | 74.166 | 0.081 | 0.550 | 0.979 |
| AID C:G % | 50.708 | 0.080 | 0.360 | 0.953 |
| ADAR Ti/Tv % | 77.908 | 0.071 | 0.386 | 0.968 |
| ADAR A:T % | 56.326 | 0.082 | 0.455 | 0.968 |
| APOBEC3G Ti/Tv % | 72.339 | 0.063 | 0.372 | 0.972 |
| APOBEC3G C:G % | 52.297 | 0.083 | 0.643 | 0.983 |
| APOBEC3B Ti/Tv % | 59.233 | 0.160 | 1.112 | 0.980 |
| APOBEC3B C:G % | 49.621 | 0.172 | 1.104 | 0.976 |

<sup>a</sup> Mean values from all saliva-1 samples (n=24, Group 1 and Group 2 subjects, 24 WES datasets)
<sup>b</sup> Average standard deviation from five replicates per volunteer (n=12, Group 1, 60 WES datasets)
<sup>c</sup> Standard deviation from all saliva-1 samples (n=24, Group 1 and Group 2 subjects, 24 WES datasets)

### Analysis of unmapped reads

The average number of unmapped reads was larger in saliva (60 WES datasets, mean=2,372,300, 98.4% mapping rate) than in blood (12 WES datasets, mean=334,182, 99.7% mapping rate), corresponding to a six fold higher unmapped rate in saliva (1.63% unmapped) compared to blood (0.27% unmapped). Overall, there is a 98.6% average mapping across all 72 samples and replicates (Supplementary Table 1). Quality statistics for unmapped reads are summarized at https://jpmam1.github.io/MultiQC/. Unmapped reads for volunteers HP_1 and HP_2 were extracted and aligned to the nr protein database. with read alignment rate to the NCBI nr database larger in saliva (41%) than in blood (33%). Reads that failed to align to NCBI nr were typically low quality. Unmapped reads derived from saliva, but not blood, were predominantly found to contain reads aligning to metagenomic species (Supplementary Figure 6).

## Discussion

AID, APOBEC3G, APOBEC3B and ADAR deaminases are implicated in 30%-40% of clinically curated SNPs in the OMIM database [13]. However, there is a paucity of research on deaminase-associated motifs in healthy subjects. Here, we have investigated deaminase-associated signatures in blood and saliva of healthy Caucasian subjects using whole exome sequencing. Our experimental design provided a framework to quantify variance in all SNVs, SNVs within the coding sequence of genes, deaminase-associated coding SNVs, and provided measures of deaminase strand-bias and transition/transversion in a highly robust and reproducible manner. Using different biological and technical sample replicates we explored differences between concordant and discordant SNV calls across the 24-subject cohort, showing strong intraclass correlation between sample replicates. No significant differences in discordant SNV calls were detected in pairwise comparisons between sample types. The sources of discordant SNVs were investigated and were found to be associated with low read depth, high strand bias, and low genotype quality. Analysis of putative somatic variants showed no conclusive evidence of somatic mutation when comparing blood and saliva samples. On average, approximately 2% of reads failed to align to the human genome, with reads derived from saliva samples primarily related to metagenomic taxa associated with the oral microbiome [41,42]. Here, we establish that saliva and blood are both appropriate sources of DNA for WES analyses, with no detected difference in ability to resolve SNVs and deaminase-associated signatures and metrics.

A key component of the experimental design in this study (Figure 1) was the capacity for comparisons between biological and technical replicates. The replicate extraction of Saliva-1 DNA, and replicate library preparation of Saliva-2, provides a measure of lab-based technical variation. Our study showed that the differences between blood and saliva, and between biological saliva replicates were very small and no larger than the level of noise of the technical replicates. Discordant SNV calls are predominantly in low coverage and/or low-quality regions of the exome. These discordant SNVs were less likely to be found in dbSNP and were dramatically enriched for known problematic SNV calls in gnomAD. These results indicate the filtering for SNVs that fail gnomAD quality analysis would improve overall reproducibility of WES SNV analysis. By filtering these failed gnomAD variants, only 3% of concordant SNVs are eliminated, but 30% of discordant SNVs are removed. These results may advise filtering strategies in future studies.

Previous research has shown high quality DNA can be obtained from saliva and blood, with results from whole exome sequencing found to be comparable for specific applications [25,26,43]. Due to oral microbiome and off-target capture, the overall unmapped rate of saliva in this study was six fold higher than that of blood (1.6% vs 0.3%), providing sufficient unmapped reads to perform a limited metagenomic analysis. Given the importance of the relationship between the microbiome and immunity [44], the oral microbiome information provided from off-target saliva exome capture may prove useful for a variety of applications in future studies (e.g. Kidd et al. [25]).

Accumulation of a small number of ‘age-related’ (pre- or non-cancerous) somatic mutations has been reported in several recent studies [16,45,46]. Despite our comprehensive analysis of WES data for evidence of somatic mutations across different ages and sexes, we were unable to unambiguously detect somatic mutations by comparing blood and saliva in these healthy individuals. Our use of biological and technical saliva replicates revealed similar numbers of candidate somatic SNVs in both technical and biological replicates for all subjects. This indicates a high level of noise and coincides with recent analyses of false-positive variant calls [47]. Bespoke somatic SNV detection approaches are evidently required to identify somatic SNVs in healthy subjects, using more advanced sequencing techniques, different cell types and more sophisticated bioinformatics [16].

There are many challenges in accurately resolving germline and somatic SNVs. Sequencing and bioinformatics artefacts are known to result in incorrect SNV calls, with numerous studies investigating the effects of exome capture kits, sequencing platform, and bioinformatics software on the ability to accurately detect SNVs [48,49]. As reported by others, performance of pipelines according to a ‘gold standard’ (such as “genome in a bottle”) does not necessarily indicate performance on ‘real world’ datasets [50]. In this study, the use of sample replicates enabled us to quantify noise and evaluate the reproducibility of SNV calls, to identify discordant germline SNVs as potential false positives and eliminate false-positive somatic SNVs. Reducing false-positive SNVs is necessary to accurately resolve deaminase-associated SNV profiles and for understanding the implications of deaminase signatures in health and disease.

Deaminase mutagenesis is associated with an increasing number of viral infections and cancer types [6,19,51–58]. With development of more advanced sequencing technologies, we will be able to detect evidence of deaminase activity with greater accuracy and examine changes over time [1,59,60]. In addition to the well-characterized effects of deaminases on genome stability in cancer, and more recently in precancerous conditions [55,61,62], deaminases have emerged as a driving factor in many human SNPs [13]. Despite the limitations of 24 individuals, and all having Caucasian ancestry, this study enabled us to investigate candidate somatic SNVs and provided us with a robust and reliable measure of deaminase-associated germline variants in healthy subjects.

## Conclusions

A large proportion of disease-associated germline variants are linked to deaminase activity. We have established that saliva and blood are appropriate sources of DNA for whole exome sequencing, with no difference in ability to resolve deaminase-associated metrics. Deaminase-associated mutations are important in pre-cancerous conditions, and in cancer, however no somatic SNVs were identified when comparing blood and saliva of healthy individuals. Investigation into the implications of deaminase activity on genome stability in healthy individuals will required more technically advanced approaches.

## Abbreviations

ADAR: <u>A</u>denosine <u>D</u>eaminase <u>A</u>cting on <u>R</u>NA;
AID: activation induced cytidine deaminase, a APOBEC family member;
APOBEC family: generic abbreviation for the deoxyribonucleic acid deaminase family (APOBECs 1,2,4 and 3A/B/C/D/F/G/H);
CDS: Coding sequence.
Deaminase: zinc-containing catalytic domain in ADAR and AID/APOBEC enzymes;
HP: Healthy Population (or Person);
ICC: Intraclass Correlation Coefficient;
R: Adenosine (A) or Guanine (G), purines;
SD: standard deviation;
SNP: single nucleotide polymorphism;
SNV: single nucleotide variant;
TSM: targeted somatic mutations;
W: weak base pair involving A or T;
Y: pyrimidines T or C.;
WES: whole exome sequencing.

## Declarations

### Ethics approval and consent to participate

This project was approved by the Monash Health Human Research Ethics Committee (16281L: “A study to measure the Targeted Somatic Mutation (TSM) test platform performance characteristics and evaluate its suitability for clinical use”). All study subjects signed written informed consent forms which were approved by the Ethics Committee.

### Consent for publication

“Not applicable”

### Availability of data and material

The data are not publicly available due to information that could compromise research participant privacy and consent. The data that support the findings of this study are available on reasonable request from the corresponding author NEH.

### Competing interests

All authors declare that they have no competing interests.

### Funding

The work was fully funded by GMDx Group Ltd (Melbourne, Australia), as a part of a GMDx Group translational research program.

### Authors’ contributions

RAL conceived the study, NEH and RAL designed the study with CF and RB. NEH and JM analyzed and interpreted the genomic data and wrote the manuscript. PR and RB were involved in implementing the clinical trial. RAL and EJS contributed to the writing of the manuscript. All authors read and approved the final manuscript.

## Acknowledgements

We thank Christopher Pendlebury and Richard Rendell from Applied Precision Medicine for TSM computational platform development and implementation. The authors also thank Trevor Wilson, Niro Pathirage, Roxane Legaie and Wishva Herath from the Monash Health Translation Precinct (MHTP) Medical Genomics Facility for exome sequencing and data processing and Margaret Smith from smartDNA for DNA extraction. Lastly, we wish to acknowledge the contributions of the study volunteers who provided samples.

## Supplementary Figure Legends

**Supplementary Figure 1: Relationships of SNV calls between and among sample replicates.**

Venn diagrams of exome SNV calls for five replicates for twelve healthy subjects. Saliva-1, and saliva-2 are replicate saliva samples. Saliva-1C and saliva-2A are technical replicates of saliva-1 and saliva-2. SNVs called in all five samples are termed *concordant* SNVs, and those not in all samples are termed *discordant* SNVs. Venn diagrams were generated online at http://bioinformatics.psb.ugent.be/webtools/Venn/.

**Supplementary Figure 2: Relationships of CDS SNV calls between and among sample replicates.**

Venn diagrams of coding sequence (CDS) SNV calls for five replicates for twelve healthy subjects.

**Supplementary Figure 3: Calculated metrics for all sample replicates**

Bar graphs of 72 exome data sets across 24 healthy individuals. Bars are colored according to the biological or technical replicate. Metrics presented are:

**SNV**, total number SNV calls,

**CDS**, number of SNVs in the coding regions,

**AID%**, percentage of CDS SNVs matching the AID deaminase motif WR<u>C</u> / <u>G</u>YW,

**ADAR%**, percentage of CDS SNVs matching the ADAR deaminase motif W<u>A</u> / <u>T</u>W, **APROBEC3G%** percentage of CDS SNVs matching the APOBEC3G deaminase motif C<u>C</u> / <u>G</u>G, and **APOBEC3B%**, percentage of CDS SNVs matching the APOBEC3B deaminase motif T<u>C</u>W / W<u>G</u>A where W=A or T, Y=C or T, R=A or G.

**Supplementary Figure 4: Depth profiles of concordant and discordant blood and saliva replicate SNVs.**

Sequencing depth of five sample types for concordant SNVs compared to discordant SNVs (n=12).

**Supplementary Figure 5: Distribution and density of candidate somatic SNVs**

Number of candidate SNVs per sample, grouped by “tumor-normal” comparison type. For ‘blood vs saliva’, blood was treated as the *tumor* sample and saliva as *normal*; for “saliva vs blood”, saliva was treated as the *tumor* sample and blood as *normal*; for “saliva biological replicates”, saliva-1 samples were compared against saliva-2 samples, and vice versa; and for “saliva technical replicates”, saliva 1 and saliva-1C samples were compared, and saliva-2 and saliva-2A samples were compared. Boxplots illustrate the median and interquartile range (IQR), with outliers defined as 1.5*IQR. The distribution for each grouping is shown above each boxplot. The number of candidate SNVs detected between technical replicates demonstrates a high level of noise across all candidate somatic SNVs. Filtering and analysis suggests all candidate SNVs are likely false positives.

**Supplementary Figure 6: Metagenomic analysis of unmapped reads**

Alignment of unmapped reads for subjects HP_1 and HP_2 to the NCBI nr protein database representing the sources of off-target WES DNA. The majority of unmapped reads derived from saliva corresponded to prokaryotic organisms associated with the oral microbiome. Unmapped reads derived from blood samples were either low quality or predominantly mapped to viral DNA (Phi-X spike-in).

