## Supplementary figures and images for "Deaminase associated single nucleotide variants in blood and saliva-derived exomes from healthy subjects"

### Supp figure 1

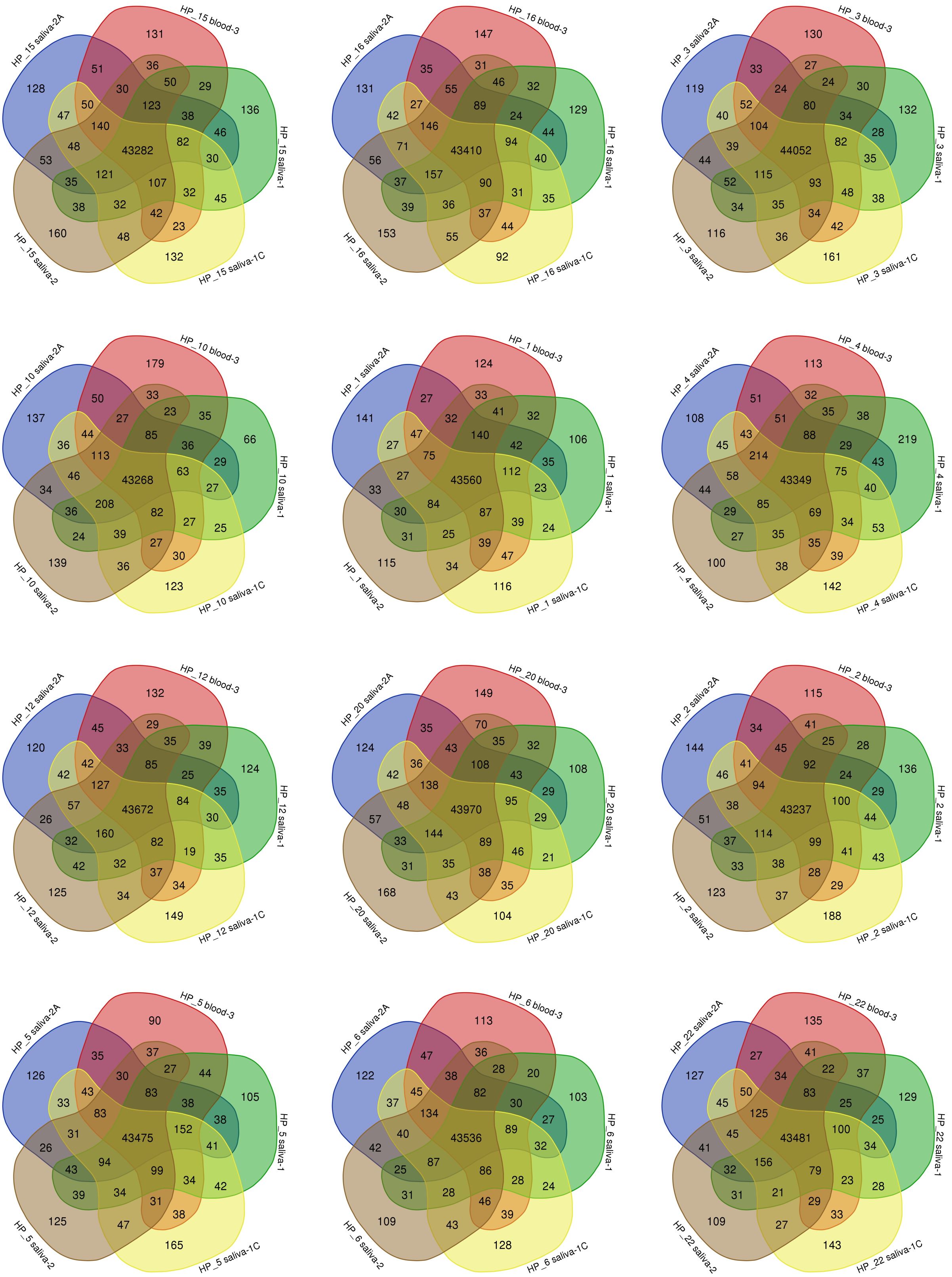

### Supp figure 2

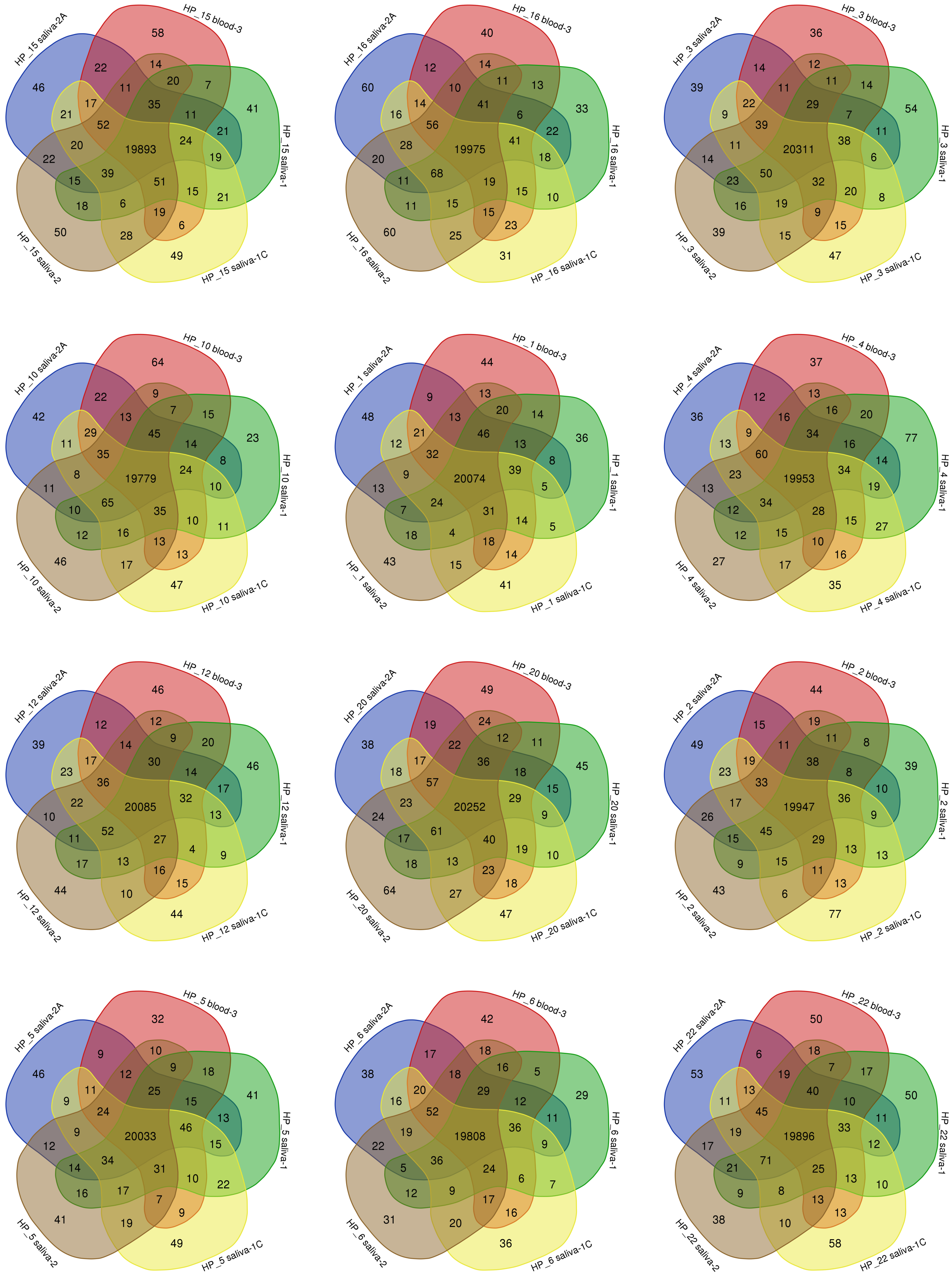

### Supp figure 3

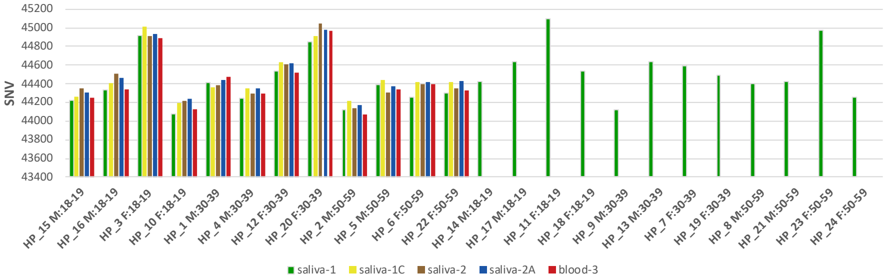

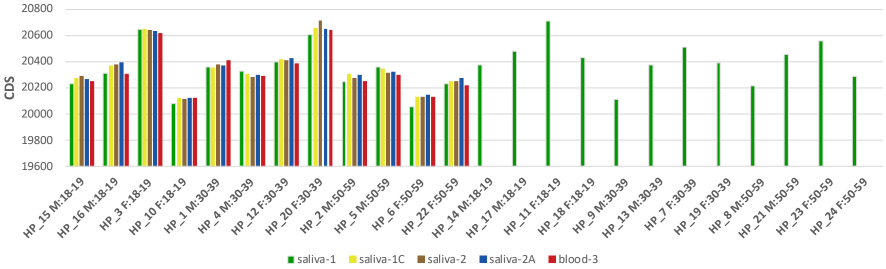

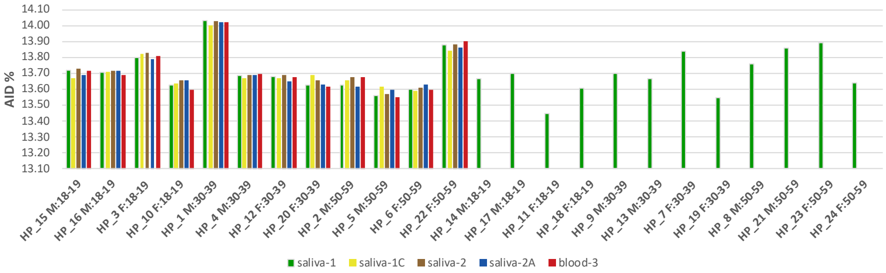

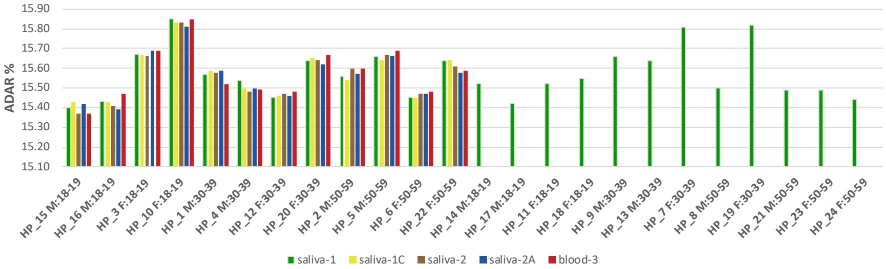

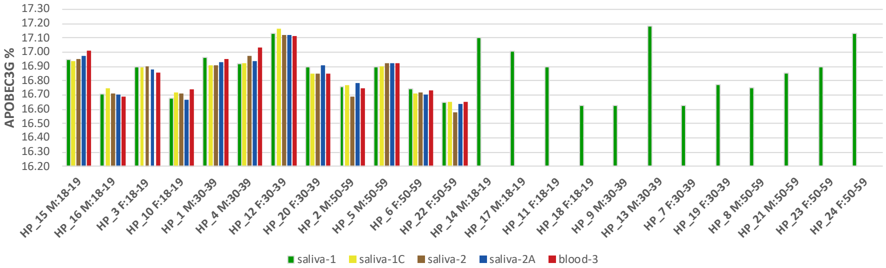

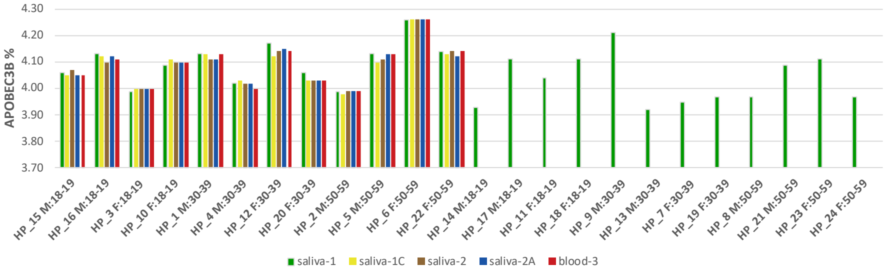

### Supp figure 4

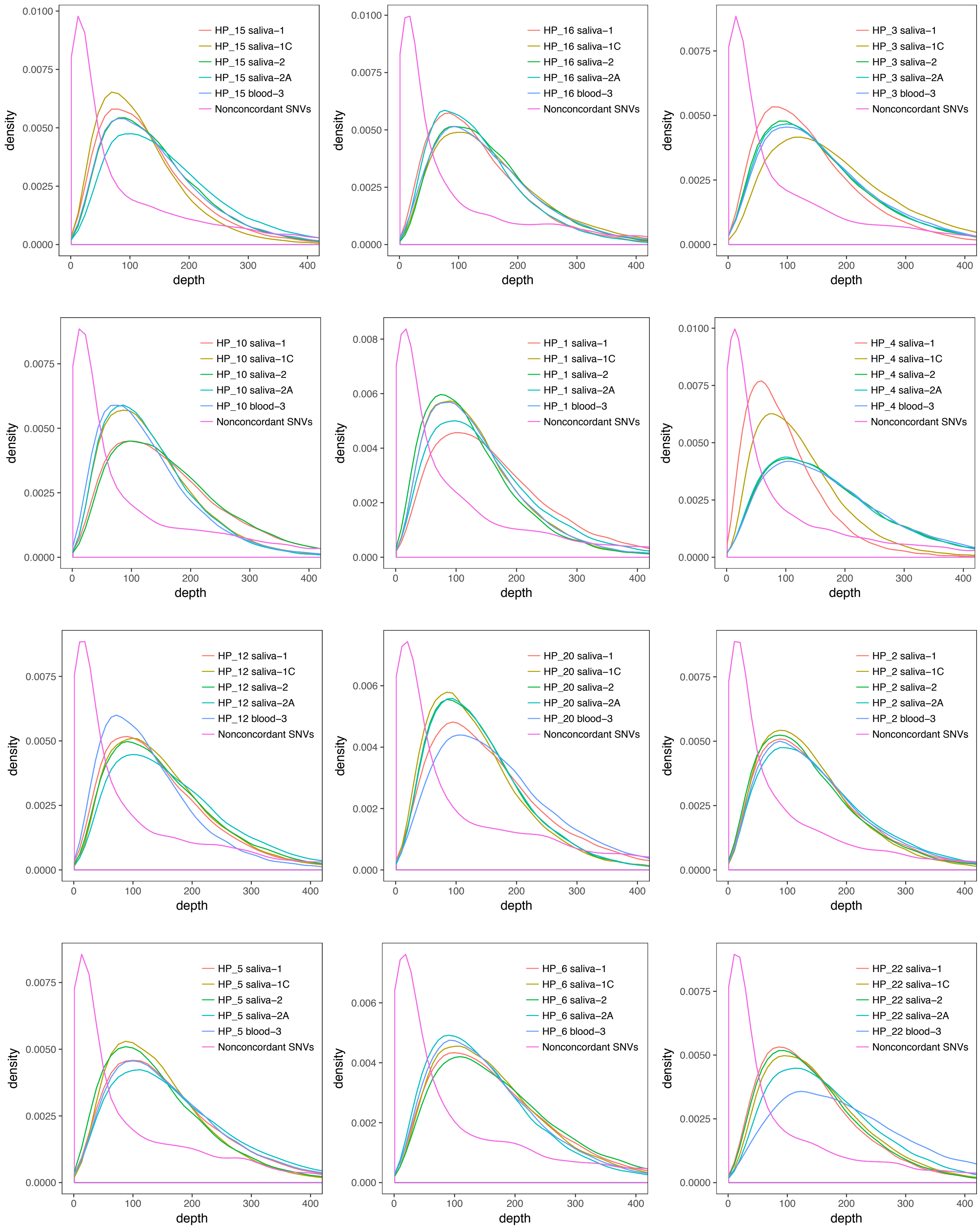

### Supp figure 5

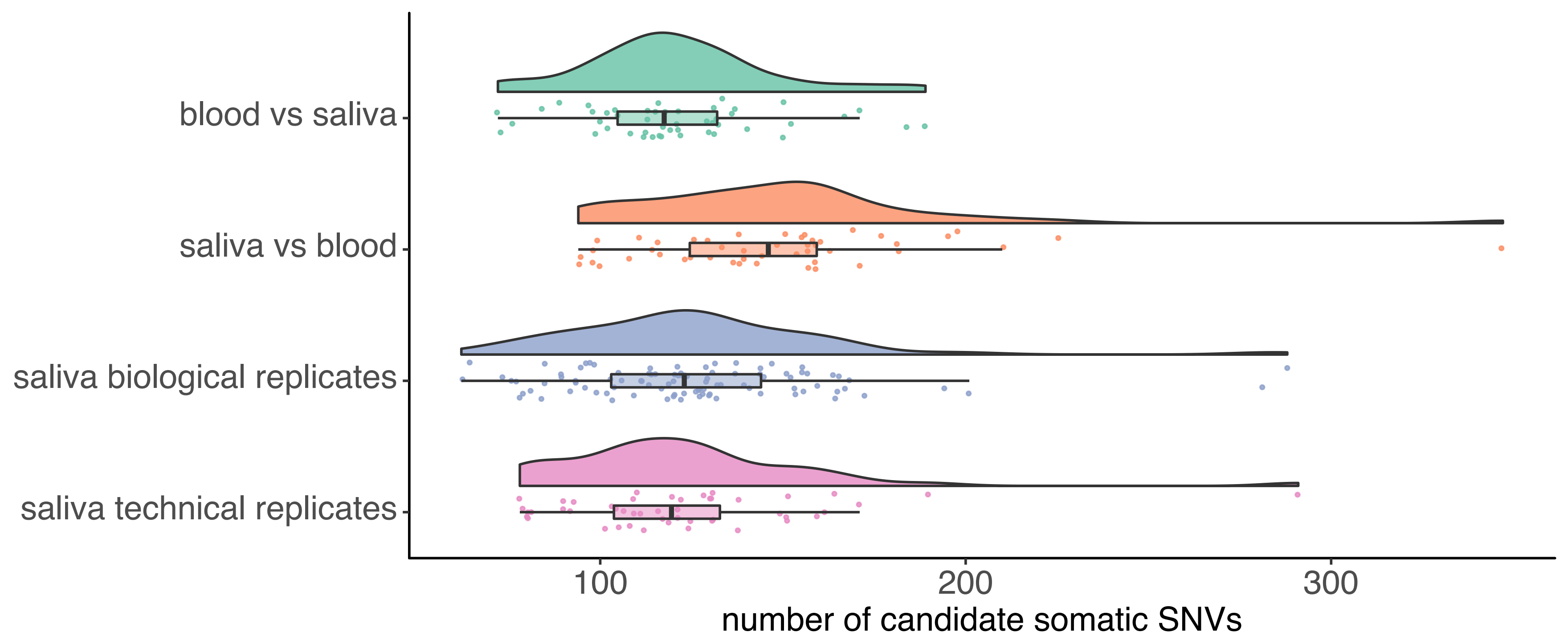

### Supp figure 6

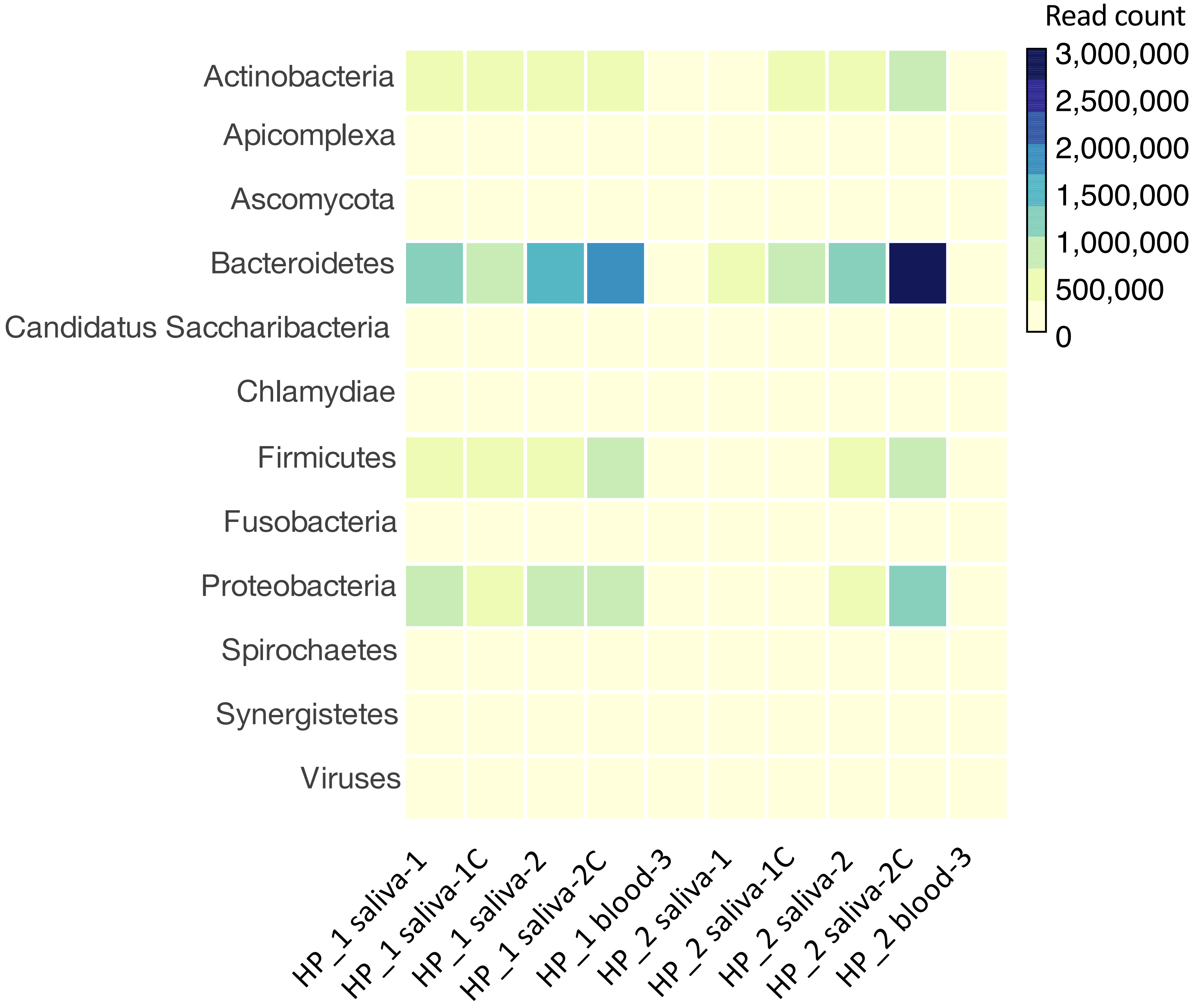
